## Supplementary Information for "A single cluster of RNA Polymerase II molecules is stably associated with active genes"

### Supplementary Figures

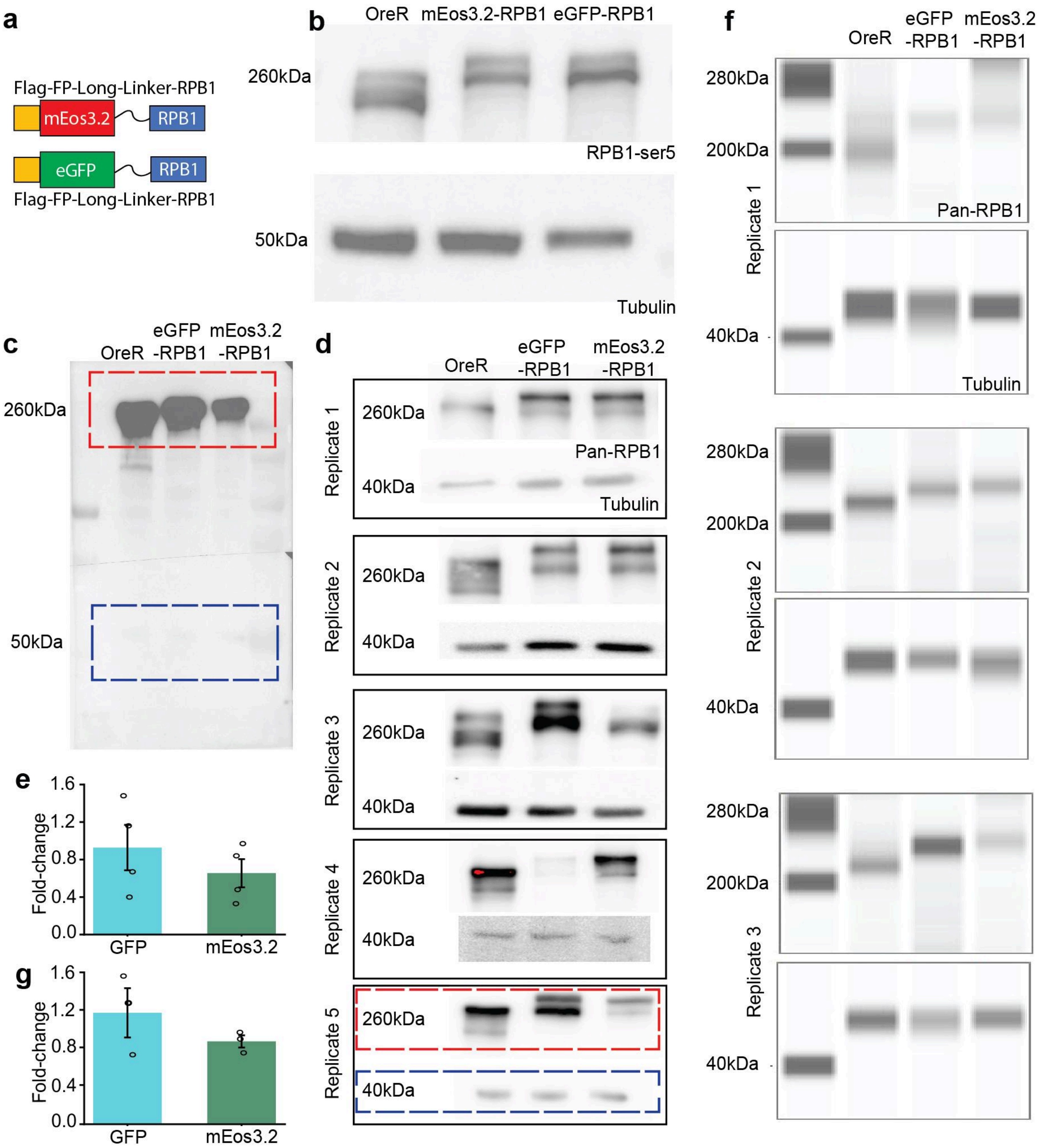


**Fig. S1.1. Constructs used for imaging. (a)** Both constructs were knocked in at the endogenous RPB1 locus using the CRISPR-Cas9 system and resulting fly lines are homozygous-viable. Each construct includes a Flag-tag at the N-terminus of the fluorescent protein (eGFP or mEos3.2) and is linked to RPB1 via a long flexible linker. **(b)** Western blots using Ser5P antibody against eGFP and mEos3.2-tagged RPB1. The double bands in RPB1 blot may correspond to previously reported phosphorylated and unphosphorylated states of the RPB1 C-terminal domain[1]. **(c)** A representative full Western blot using pan-RPB1 and alpha-Tubulin antibodies. An exposure time of 30 sec was used here so that the control and RPB1 are visible in the same image. **(d)** Zoomed in regions for the RPB1 and corresponding Tubulin bands for all the pan-RPB1 antibody replicates. Dashed red and blue rectangles represent the replicate that corresponds to the rectangles shown in the full blot. Exposure times of 5, 10, 15, 30, and 30 sec were used for replicates 1, 2, 3, 4, and 5 respectively. **(e)**

Quantification of protein concentration normalized to the OreR across the Western blot replicates for the pan-RPB1 antibody antibody shown in (d). Error bars represent standard deviation. No significant difference was measured between eGFP and mEos3.2 ($p = 0.458$ from paired two-sided t-test). **(f)** RPB1 and Tubulin bands for three replicates conducted on the Jess automated Western blot system as an orthogonal measurement due to variability discovered in the manually conducted blots in (g). All blots were imaged using the default High Dynamic Range 4.0 feature. **(g)** Quantification of protein concentration normalized to OreR across replicates for the eGFP and mEos3.2 constructs obtained from the Jess system shown in (f). No significant difference was measured between eGFP and mEos3.2 ($p = 0.416$ from paired two-sided t-test). Source data are provided as a Source Data file.

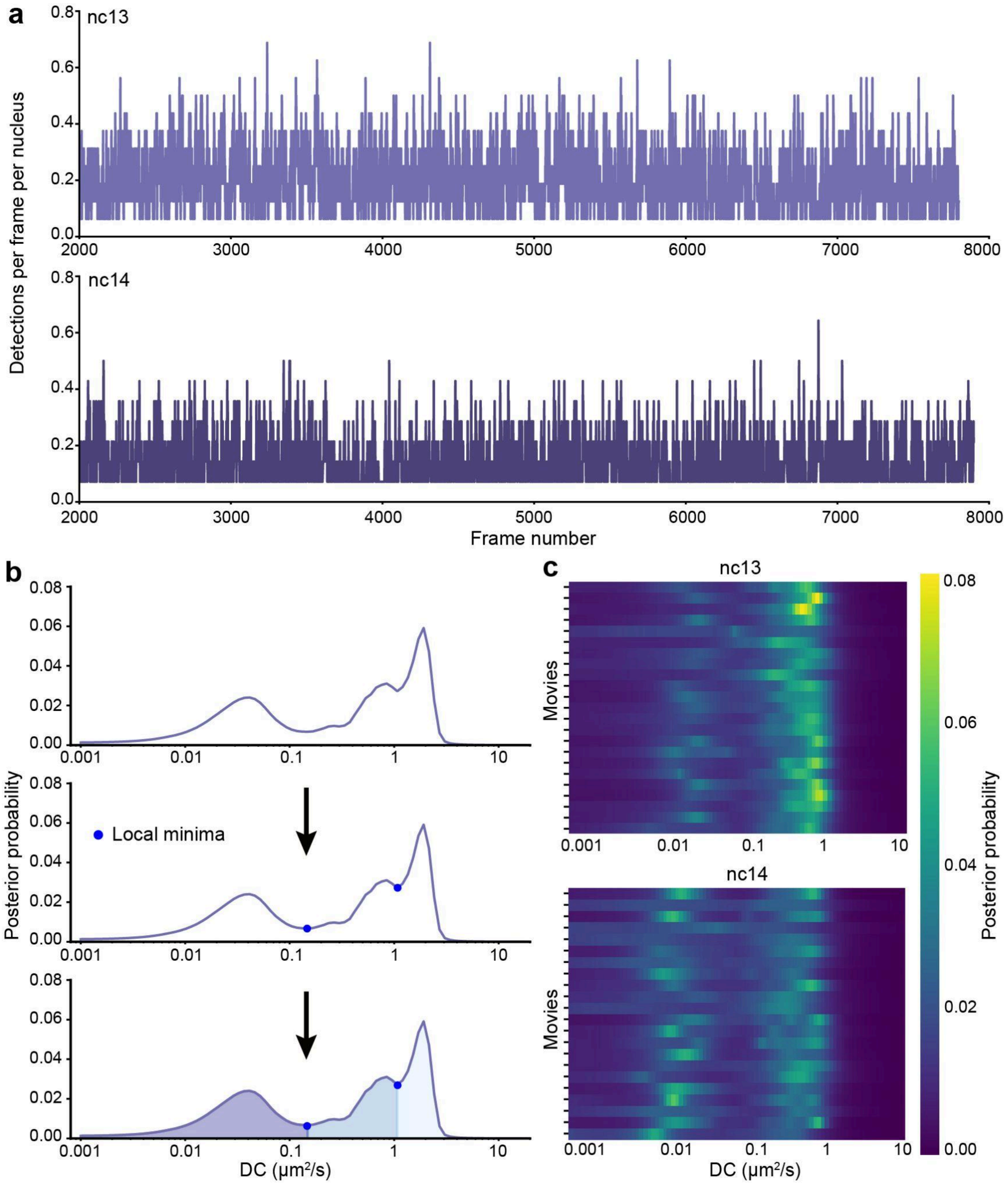


**Fig. S1.2: Single-molecule tracking quality control. (a)** Representative traces showing the number of detections per frame per nucleus in nc13 (top) and nc14 (bottom). **(b)** Analysis pipeline showcasing how the diffusion coefficient spectrum is divided into different kinetic states based on the local minima values. Local minima are represented by blue dots. The different kinetic states, free, intermediate and fast, are shown as differentially shaded regions. **(c)** Heatmaps of the probability distributions of diffusion coefficients obtained for each movie (one field of view/row; 8-15 nuclei per field of view) for nc13 (top) and nc14 (bottom).

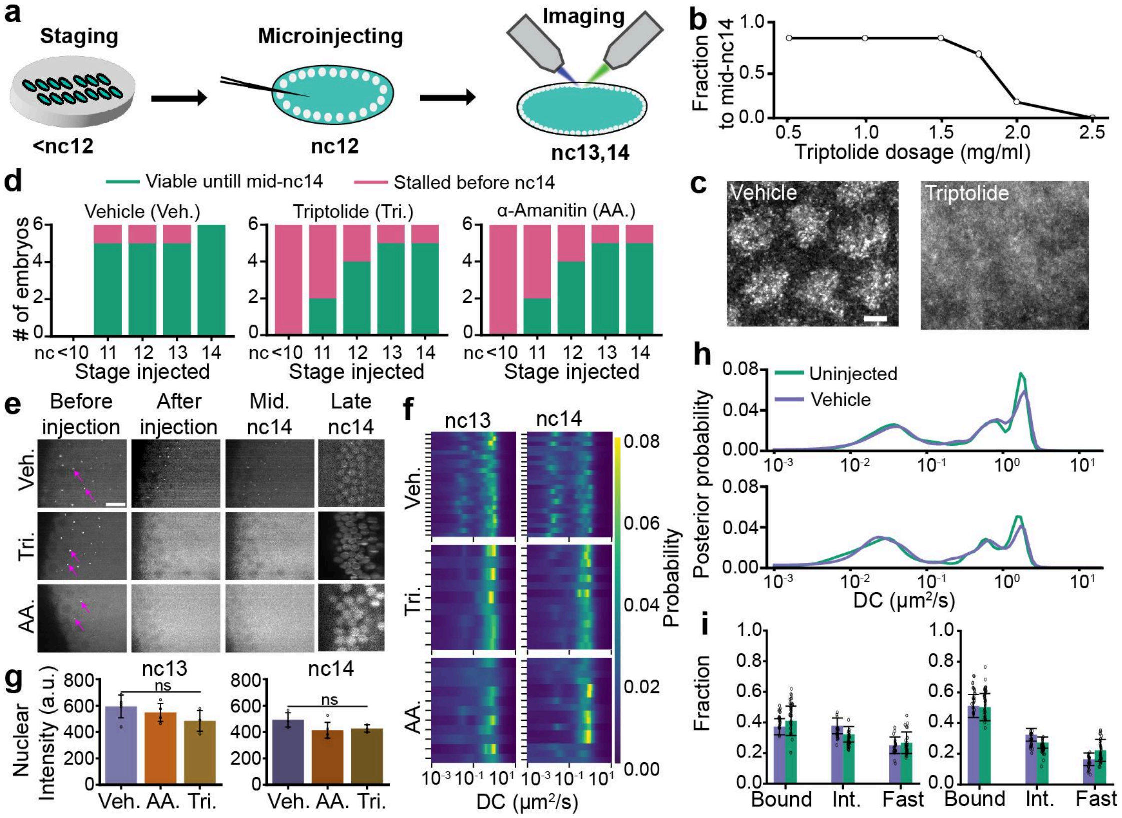


**Fig. S2.1: Validation of the microinjection process. (a)** Schematic showing an overview of the different stages of the microinjection process. **(b)** Triptolide dosage curve ranging from 0.5 mg/ml to 2.5 mg/ml. A minimum of 6 embryos were considered for each dosage. **(c)** Representative images of nc14 embryos injected with EUTP (left) and EUTP + triptolide (right). A 0.5 mg/ml concentration of triptolide was used for these injections. The scale bar is 2 μm. **(d)** Quantification of the microinjection survival assay for vehicle injections (left), triptolide injections (center) and α-amanitin injections (right). **(e)** Representative images of embryos prior to injection showing *hunchback* MS2 signal (example spots indicated by pink arrows). Injection with either triptolide or α-amanitin causes the MS2 spots to disappear but after injecting with the vehicle the MS2 spots remain. Late in nc14 (~45 minutes into nc14) the embryo injected with the vehicle continues to develop normally while both of the drug injected embryos show major developmental defects validating the injection process. **(f)** Heatmaps of the probability distributions of diffusion coefficients obtained for the vehicle injection, triptolide injection and α-amanitin injection in nc13 (left column) and nc14 (right column). Each row represents one movie. **(g)** Raw nuclear mean intensity of RNAPII. Data points are individual embryos. For vehicle-injected embryos, n=5 in nuclear cycle 13 and n=3 in nuclear cycle 14. For triptolide- and α-amanitin-injected embryos, n=3 in each nuclear cycle. Error bars represent standard deviation.**(h)** Diffusion coefficient distributions for the uninjected embryo compared to the vehicle embryo in nc13 (top) and nc14 (bottom). A total of 210,638 trajectories across 9 independent embryos and 205,532 trajectories across 10 independent embryos were obtained for nc13 and nc14 respectively for the vehicle case. A total of 203,697 trajectories across 10 independent embryos and 254,957 trajectories across 10 independent embryos were obtained for nc13 and nc14 respectively for the uninjected case. **(i)** Bound, intermediate, and fast fractions for both uninjected and vehicle in nc13 (top) and nc14 (bottom). Individual data points represent individual fields of view. Bars represent mean across the fields of view and error bars represent standard deviation. Two-sided Mann-Whitney U-test was performed to determine significance and following p-values were used: *p<0.05, **p<0.01, and ***p<0.001. No significance bars here indicate lack of statistical significance for each kinetic state between uninjected and vehicle cases. Source data are provided as a Source Data file.

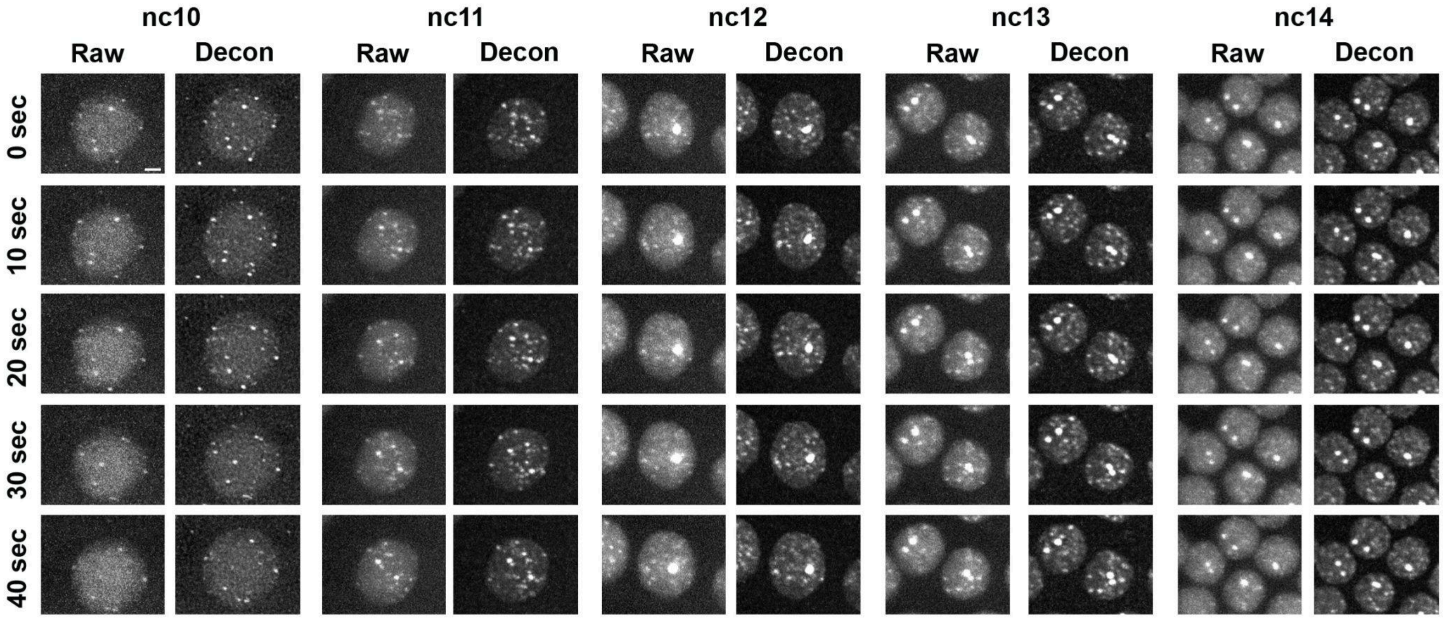


**Fig. S3.1: RNAPII clusters are visible from nc10 to nc14.** Time-lapse snapshots of eGFP-RPB1 nuclei showing five consecutive imaging time points (rows) during interphase for each nuclear cycle from nc10 to nc14 (columns). Image contrasts are all scaled equally. Images are maximum intensity projections of both the raw and corresponding deconvolved images over 16.8 µm. Scale bar is 1 µm.

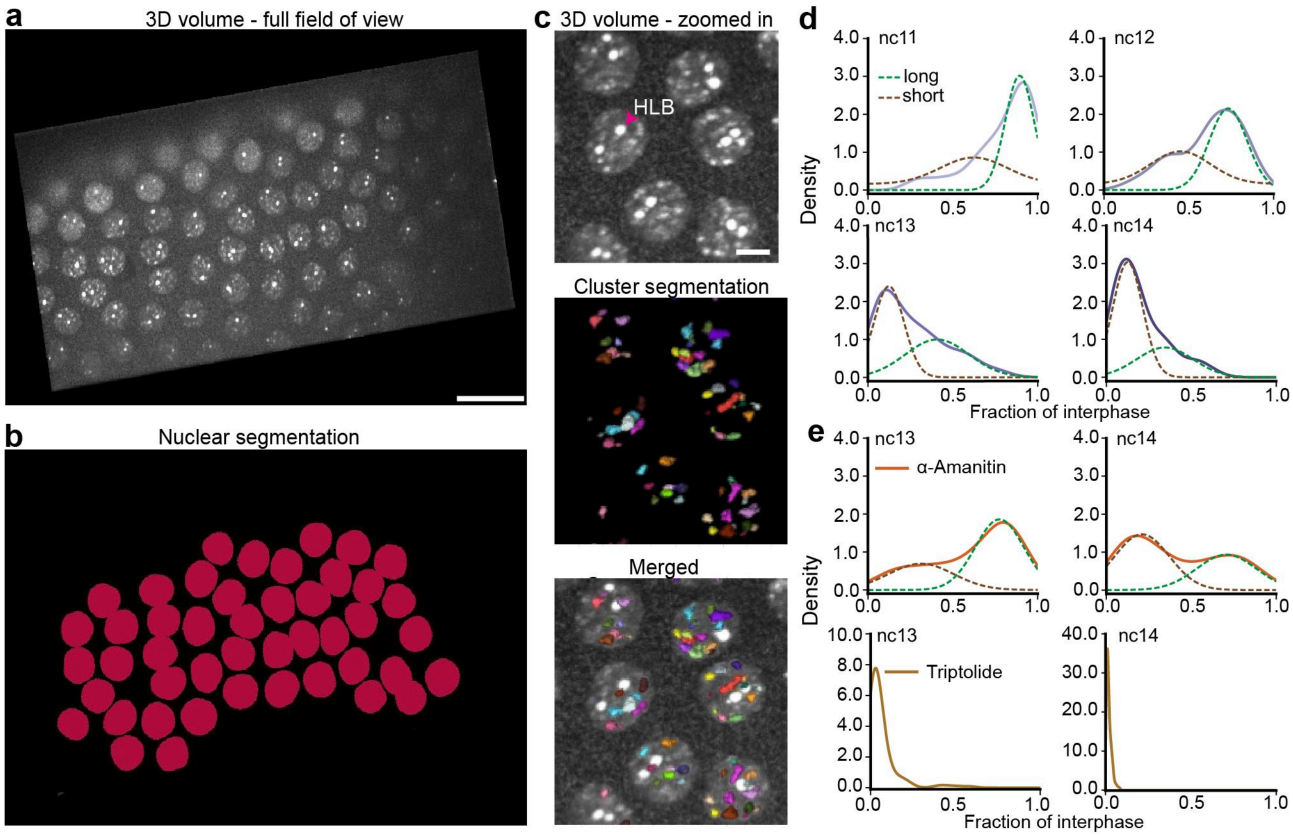


**Fig. S3.2: Characterization of RNAPII clusters. (a)** Volumes of a layer of nuclei on the *Drosophila* embryo surface are acquired during imaging and deconvolved. The scale bar is 12 μm. **(b)** Nuclei are segmented in 3D using a trained machine learning model. **(c)** Zoomed-in region of the embryo showing (top) few nuclei and RNAPII clusters, (middle) segmented clusters and (bottom) merged image. A representative Histone Locus Body (HLB) is marked by a pink arrow. The HLBs are excluded from the segmentation. The scale bar is 2 μm. Cluster segmentation was performed in 3D using a custom analysis pipeline (see Methods for more detail). **(d)** Fitting the probability density functions (PDFs) for the cluster lifetimes as a fraction of interphase length for the vehicle embryos in nc11-14. Dashed lines on each pdf represent the different gaussian fits. **(e)** Fitting the PDFs for the cluster lifetimes as a fraction of interphase length for the α-amanitin-injected embryos in nc13 (left) and nc14 (right). Dashed grey lines on each PDF represent the different Gaussian fits. Cluster lifetimes for triptolide-injected embryos could not be fit to a multi-Gaussian model, as only short-lived populations were observed. Source data are provided as a Source Data file.

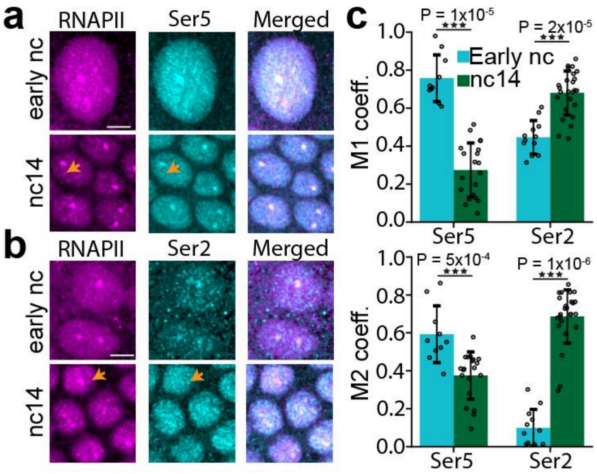


**Fig. S3.3: Immunofluorescence analysis of Ser5 and Ser2. (a)** Representative nuclei stained for eGFP-RPB1 (magenta), Ser5 (cyan), and merged channels during early nuclear cycles (nc11-12) and nuclear cycle 14. **(b)** Representative nuclei stained for eGFP-RPB1 (magenta), Ser5 (cyan), and merged channels during early nuclear cycles (nc11-12) and nuclear cycle 14. Scale bars are 4 μm. Yellow arrows in (a) show the lack of a Ser5 cluster in a corresponding RNAPII cluster. Yellow arrows in (b) show a Ser2 cluster colocalized with a corresponding RNAPII cluster. **(c)** Manders coefficient analysis of eGFP-RPB1 with either Ser5 or Ser2 channels in early nuclear cycles versus nc14 embryos. Data points show individual nuclei and error bars represent standard deviation of mean. Bars represent means across individual nuclei. Sample sizes: early nc, n=10 (Ser5), n=12 (Ser2); nc14, n=20 (Ser5), n=25 (Ser2). Data collected from a minimum of two embryos per condition. Two-sided Mann-Whitney U-test was performed to determine significance and following p-values were used: *p<0.05, **p<0.01, and ***p<0.001. Source data are provided as a Source Data file.

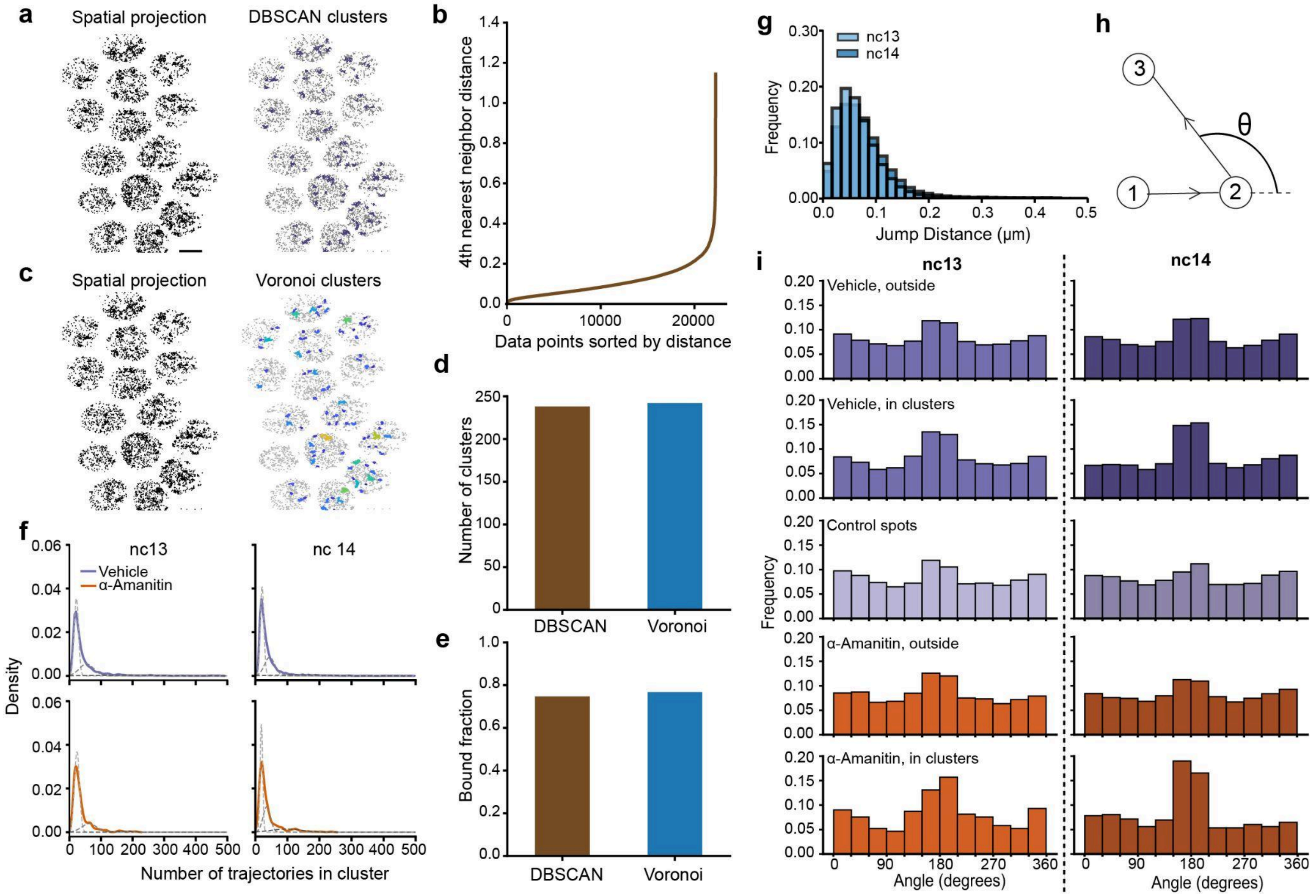


**Fig. S4.1**: **Cluster calling validation and anisotropy calculation. (a)** Representative spatial mapping of average single molecule trajectories (left) and the clusters that are identified on the same field of view using DBSCAN. Scale bar is 4 μm. **(b)** Representative elbow curve analysis to determine DBSCAN parameters. **(c)** Clusters were identified using the Voronoi tessellation method as an orthogonal approach to validate DBSCAN-based cluster detection. **(d)** Number of clusters identified using the two different cluster calling methods in the same embryo and same field of view. **(e)** RPB1 bound fraction inside the clusters determined by two different cluster calling methods in the same embryo and same field of view. **(f)** Probability density functions for the number of trajectories inside clusters in nc13 (left) and nc14 (right) for vehicle embryos (top row) and α-amanitin injected embryos (bottom row). Dashed lines show the 3-state fits for each condition. Number of clusters are 1,517 and 202 for vehicle and α-amanitin injected embryos respectively in nc13 and 1,667 and 258 for the same conditions in nc14. **(g)** Histogram of jump distances from H2B trajectories in nc13 and nc14 used to determine the distance threshold for anisotropy analysis. A total of 51,158 tracks were considered for nc14 and 39,492 for nc13. **(h)** Schematic showing which angle is considered for the calculation for every pair of displacements. **(i)** Histograms showing angle distributions for different conditions in nc13 (left) and nc14 (right). Each row represents a different condition and the condition is labelled on the top of each plot in the left column. Number of angles in the vehicle case are 13,107, 3,070, and 5,754 for outside clusters, inside clusters and in control spots respectively in nc13 and 10,540, 2,057, and 6,104 for the same categories in nc14. Number of angles in the α-amanitin injected embryos are 2,664, 344 and 2,189 for outside clusters, inside clusters, and in control spots respectively in nc13 and 4,807, 448, and 2,894 for the same categories in nc14. Source data are provided as a Source Data file.

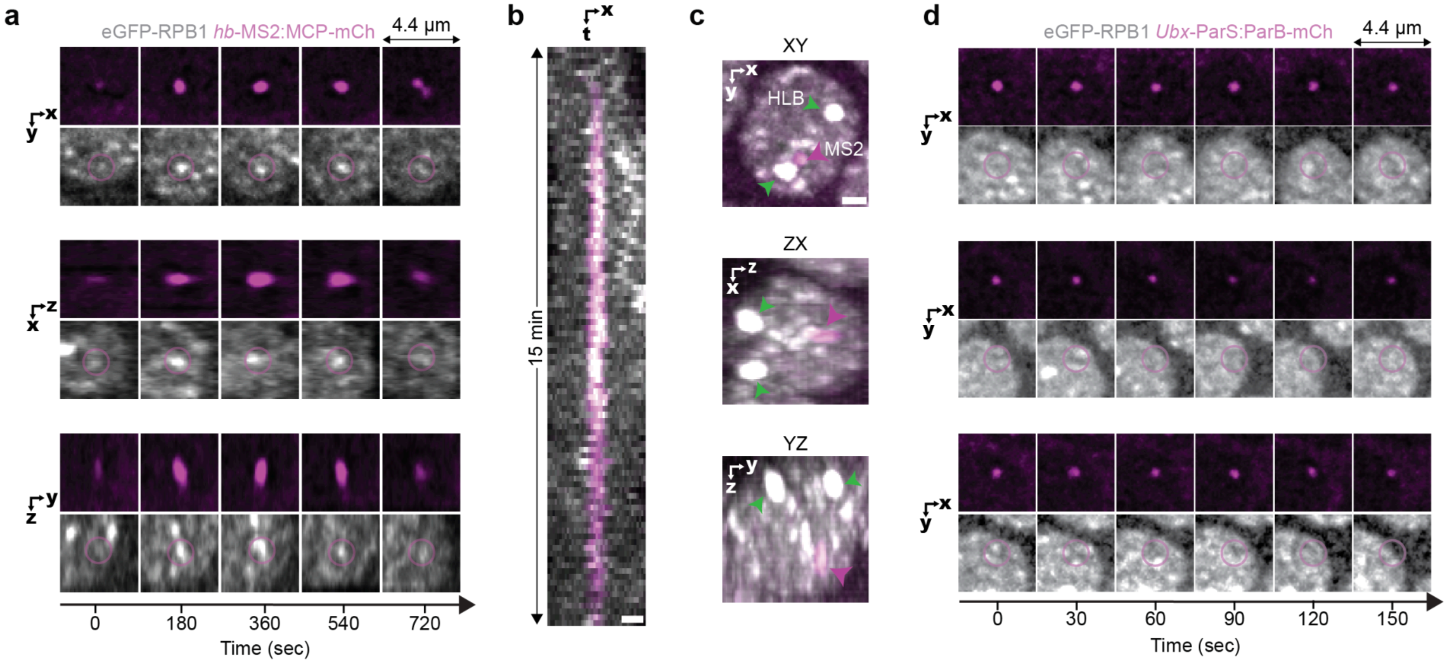


**Fig. S5.1**: **Measurement of RNAPII and MCP intensities around MS2 spots. (a)** Orthogonal view images of a nucleus in nc14 with eGFP-RPB1 (grey) and MCP-mCherry (pink) labelling active transcription at a *hb*-MS2 reporter during a ~15 min transcriptional burst. Cytoplasmic MCP signal is present as it is expressed without a nuclear localization signal. **(b)** Kymographs for the same nucleus calculated as max-projection over a 1.2 µm y-slice. The scale bar is 1 µm. **(c)** Orthogonal view max projections for an example nucleus at a time point where the *hb*-MS2 reporter is actively transcribing, showing that the MS2 spot (magenta arrow) is not associated with HLBs (green arrows). Intensities for each orthogonal view were scaled independently. The scale bar is 1 µm. **(d)** Example nuclei in nc14 with eGFP-RPB1 (grey) and ParB-mCherry (pink) labelling an inactive *Ubx*-ParS2 construct. Nuclei were imaged every 30 sec to allow for longer observation times of the ParB-mCherry signal.

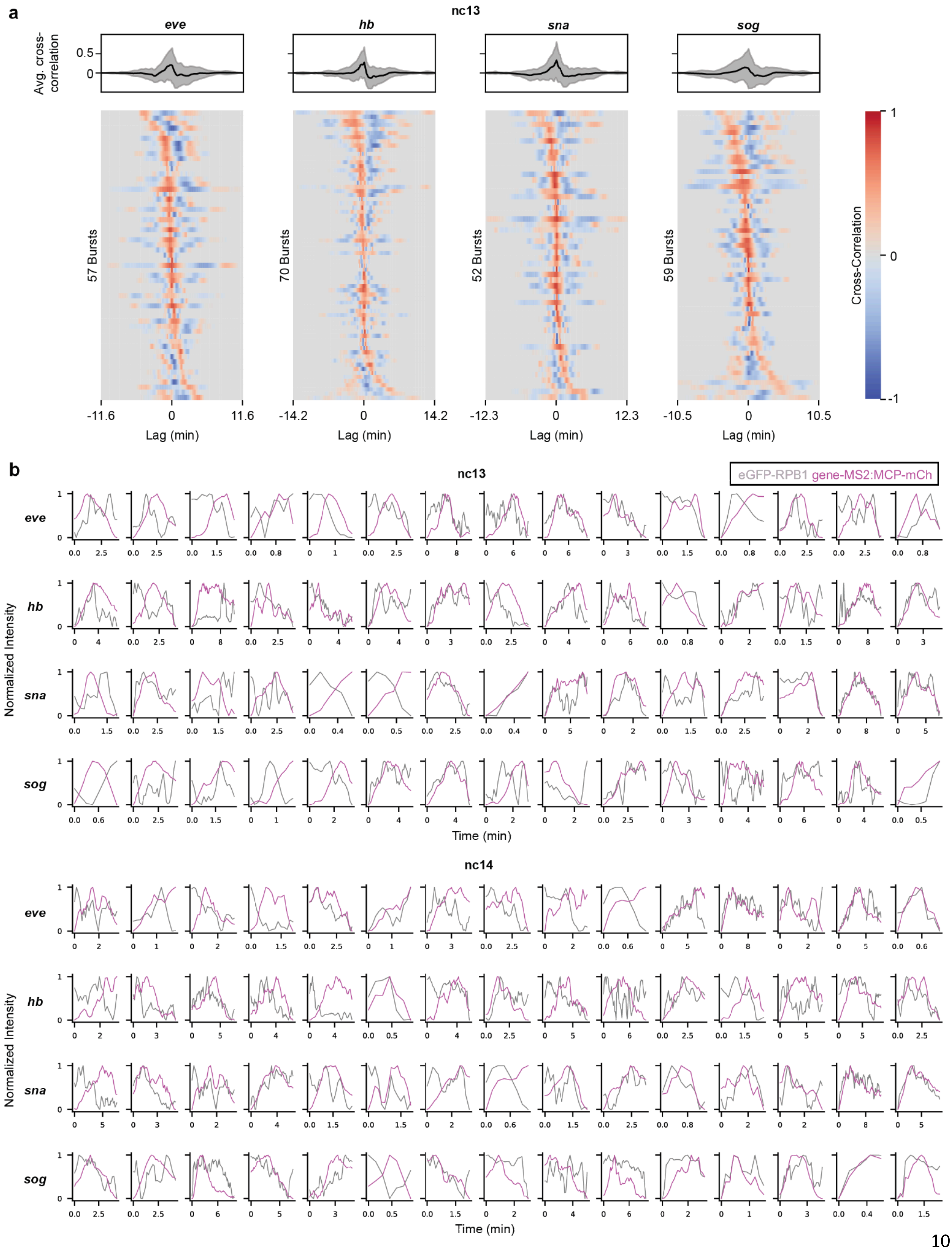
a
nc13
eve
hb
sna
sog
Avg. cross-correlation
57 Bursts
70 Bursts
52 Bursts
59 Bursts
Cross-Correlation
Lag (min)
b
nc13
eGFP-RPB1 gene-MS2:MCP-mCh
Normalized Intensity
Time (min)
nc14

**Fig. S5.2: RNAPII and MCP intensities are highly correlated at active genes. (a)** Average cross-correlation (top) and individual cross-correlations (bottom) of eGFP-RPB1 and MCP-mCh intensities in nuclei in nc13 from 3 biological replicates. Gray shaded area in line plots represent standard deviation. RPB1 intensity was normalized to reflect enrichment above nuclear background. **(b)** Representative traces of RPB1 and MCP at each of the 4 reporter genes in nc13 and nc14.

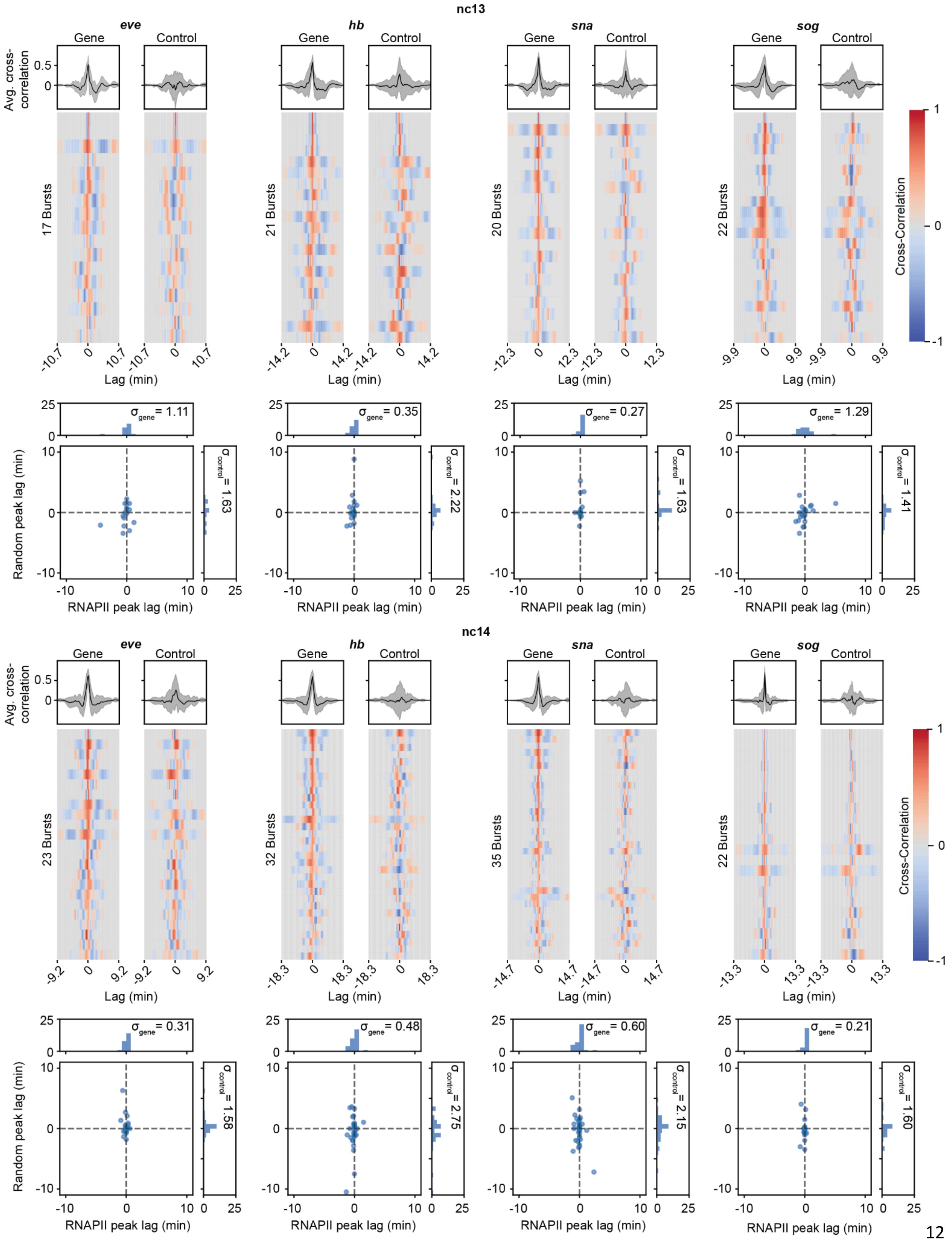

nc13
eve
hb
sna
sog
Gene
Control
Avg. cross-correlation
17 Bursts
21 Bursts
20 Bursts
22 Bursts
Lag (min)
Cross-Correlation
σgene = 1.11
σgene = 0.35
σgene = 0.27
σgene = 1.29
σcontrol = 1.63
σcontrol = 2.22
σcontrol = 1.63
σcontrol = 1.41
Random peak lag (min)
RNAPII peak lag (min)
nc14
23 Bursts
32 Bursts
35 Bursts
22 Bursts
σgene = 0.31
σgene = 0.48
σgene = 0.60
σgene = 0.21
σcontrol = 1.58
σcontrol = 2.75
σcontrol = 2.15
σcontrol = 1.60

**Fig. S5.3: RNAPII intensities at control spots are not correlated with MCP intensity at active genes.** Heatmaps show correlations for a subset of highly-correlated eGFP-RPB1 and MS2:MCP-mCh traces (left), and correlations between eGFP-RPB1 intensities in control spots and MS2:MCP-mCh traces for the same nuclei (right) for each reporter gene in nc13 (top) and nc14 (bottom). Scatter plots and histograms show the spread of peak lag times in each pair of cross-correlations.

### References:

1. Anti-RNA polymerase II RPB1 (phospho S5) antibody. https://www.abcam.com/en-us/products/primary-antibodies/rna-polymerase-ii-rpb1-phospho-s5-antibody-ab240740#.
